# WSInsight: a cloud-native, agent-callable platform for single-cell whole-slide pathology

**DOI:** 10.64898/2025.12.07.692260

**Authors:** Chao-Hui Huang, Oluwamayowa E. Awosika, Diane Fernandez

## Abstract

WSInsight is an open-source platform for cohort-scale H&E whole-slide image analysis. It performs patch-level inference together with single-cell segmentation and phenotype classification, with morphology- and transcriptome-supervised cell-type heads retrainable from public data. Slides are read from cloud repositories and per-slide outputs are written to QuPath and OMERO. The same workflow is AI-agent callable through a Model Context Protocol endpoint. Applying it to TCGA cohorts recovered known immune and molecular associations.

---

The digitization of glass slides has created opportunities for deep-learning-based tissue analysis, yet most existing open-source whole-slide image (WSI) analysis frameworks remain confined to local workflows that are difficult to deploy across institutions or to drive from automated systems [1–3]. Tools such as WSInfer [4], TIA Toolbox [5], SlideFlow [6], MONAI Pathology [7] and PHARAOH [8] have lowered technical barriers, while standalone single-cell models such as CellViT [9] and HoVer-Net [10] provide accurate nuclei segmentation and typing as research codebases. In practice these capabilities remain separated: patch classifiers, single-cell models, slide storage, pathology viewers, and downstream statistical workflows are typically connected by project-specific scripts, which makes cohort-scale studies difficult to reproduce and difficult to run in institutional or cloud-hosted environments (Table 1).

**Table 1:** Feature comparison of representative open-source whole-slide pathology frameworks. ✓ = native first-class support; *∼*= partial / requires user code; – = not available. Definitions of each capability column and the threshold for ✓ *∼*vs. vs. – are stated in the paragraph immediately preceding this table. All listed frame-works are open source (Apache-2.0, BSD-3-Clause, GPL-3.0 or MIT). Among the surveyed frameworks, WSInsight is the only one that satisfies the Agent-callable criterion through both routes (manifest and MCP-conformant tool surface).

| Framework | Patch-level | Single-cell | Cloud I/O | QuPath import | OMERO import | Spatial analysis | Agent-callable |
| --- | --- | --- | --- | --- | --- | --- | --- |
| WSInfer [4] | ✓ | – | ~ | ~ | – | – | – |
| TIA Toolbox [5] | ✓ | ~ | – | – | – | ~ | – |
| SlideFlow [6] | ✓ | – | – | – | – | – | – |
| MONAI Pathology [7] | ✓ | ~ | – | – | – | – | – |
| PHARAOH [8] | ✓ | – | ~ | – | – | – | – |
| CellViT [9] | – | ✓ | – | – | – | – | – |
| HoVer-Net [10] | – | ✓ | – | – | – | – | – |
| SPARK [15] | ✓ | ✓ | – | – | – | ✓ | N/A<sup>‡</sup> |
| **WSInsight (this work)** | ✓ | ✓ | ✓ | ✓ | ✓ | ✓ | ✓ |

<sup>†</sup>For the Agent-callable column, agent-readable prose documentation (e.g. README.md, AGENTS.md, SKILL.md) is not counted as a typed tool interface.
<sup>‡</sup>SPARK is a closed agentic discovery framework: it is not externally callable by other agents, but internally it orchestrates its own agents to generate and verify biological hypotheses.

WSInsight (Fig. 1) is built around three layers. At the model layer it inherits the Model-Zoo philosophy of WSInfer [4]—every existing WSInfer patch-level head loads unchanged—and extends that patch-classifier architecture with end-to-end single-cell nuclei segmentation and phenotype classification. The single-cell layer is model-agnostic: representative single-cell models including CellViT [9], HoVer-Net [10], and StarDist–ResNet50 [11] all plug into the same downstream pipeline through a common per-cell record contract.

**Fig. 1:**
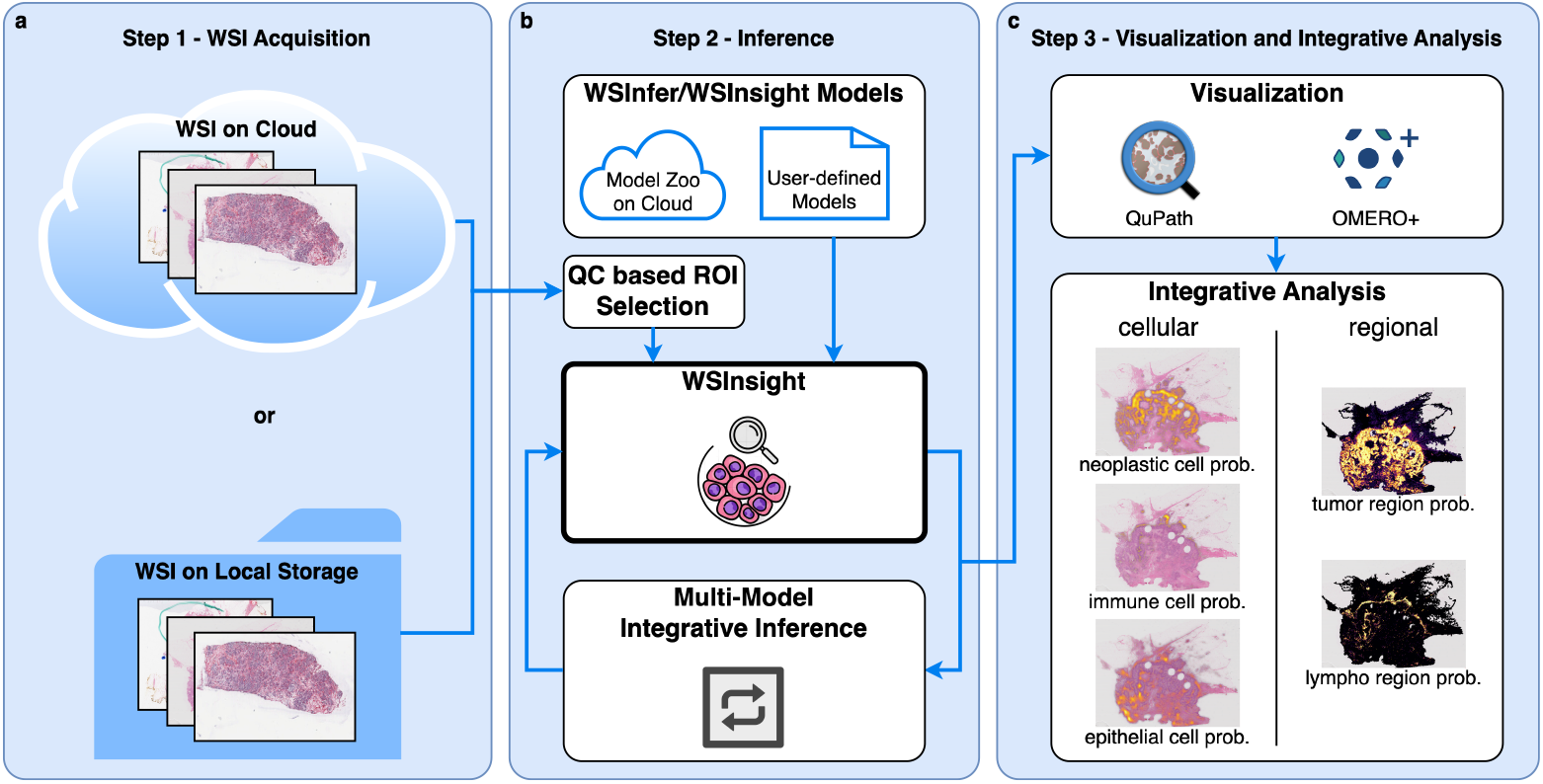
WSInsight provides an end-to-end, cloud-native workflow for whole-slide image analysis. **(a)** WSIs are accessed from cloud or local storage. **(b)** A QC-based ROI generator identifies tissue regions, and WSInsight performs patch-level classification and single-cell prediction using models from the WSInfer/WSInsight Model Zoo. **(c)** Outputs (region-level probability maps and cellular predictions) are auto-imported into QuPath and OMERO for integrated tissue interpretation.

At the I/O and viewer layer, whole-slide images are streamed on demand from local disk, S3 object storage, or the NCI Genomic Data Commons (GDC) [12], and per-slide results are returned as GeoJSON detections viewable in QuPath [13] and as OME-CSV cell tables registerable in OMERO [14] through a bundled QuPath extension and a companion OMERO module.

At the automation layer, the same command-line entry point is exposed to AI agents through a standards-conformant Model Context Protocol (MCP) interface (Fig. 2), so a workflow designed interactively in QuPath can be re-executed by an agent on a different cohort, and analyses generated by automated workflows can be rerun by human users through the same interface.

**Fig. 2:**
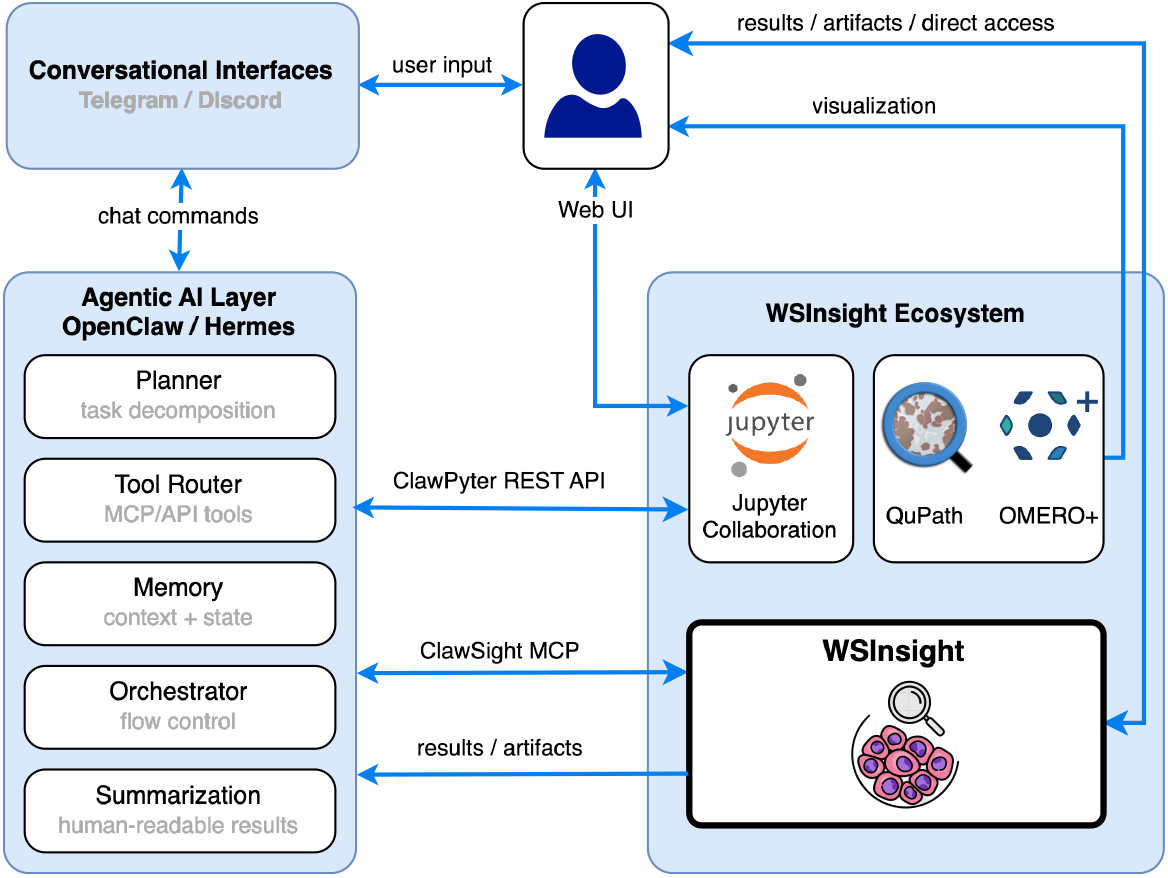
Agent-callable interface to WSInsight. WSInsight is invoked from a web UI, a JupyterLab notebook, or a chat client (Telegram, Discord) through two entry points: the ClawSight Model Context Protocol (MCP) server for AI agents, and the ClawPyter notebook bridge that lets an agent read, write, and execute Jupyter cells in the same workspace a human analyst would use. Requests can be routed through an agentic layer (OpenClaw / Hermes) that splits each request into a Planner, a Tool Router over MCP-exposed tools, a Memory module, an Orchestrator, and a final Summarization step. Per-slide outputs (probability maps, single-cell detections, and derived features) are returned to QuPath and OMERO+ for review.

To demonstrate end-to-end pipeline at The Cancer Genome Atlas (TCGA) scale, WSInsight was used to compute a unified tumour-infiltrating lymphocyte (TIL) and neighborhood-composition landscape across TCGA-BRCA (breast cancer) and TCGA-CRC (colorectal cancer), streamed directly from the GDC in one cohort-scale run (Fig. 4). These analyses are presented as a platform use case rather than as an independent biomarker-discovery study: their role is to show that the same workflow yields associations consistent with known immune and molecular patterns on whole cohorts. The run produced QuPath-ready detections (Fig. 3) and OMERO-ready cell tables for every slide; the resulting features showed associations consistent with known biology for PAM50 / receptor status in BRCA, for consensus molecular subtype (CMS) / microsatellite-instability (MSI) status in CRC, and for immune-microenvironment readouts in both indications.

**Fig. 3:**
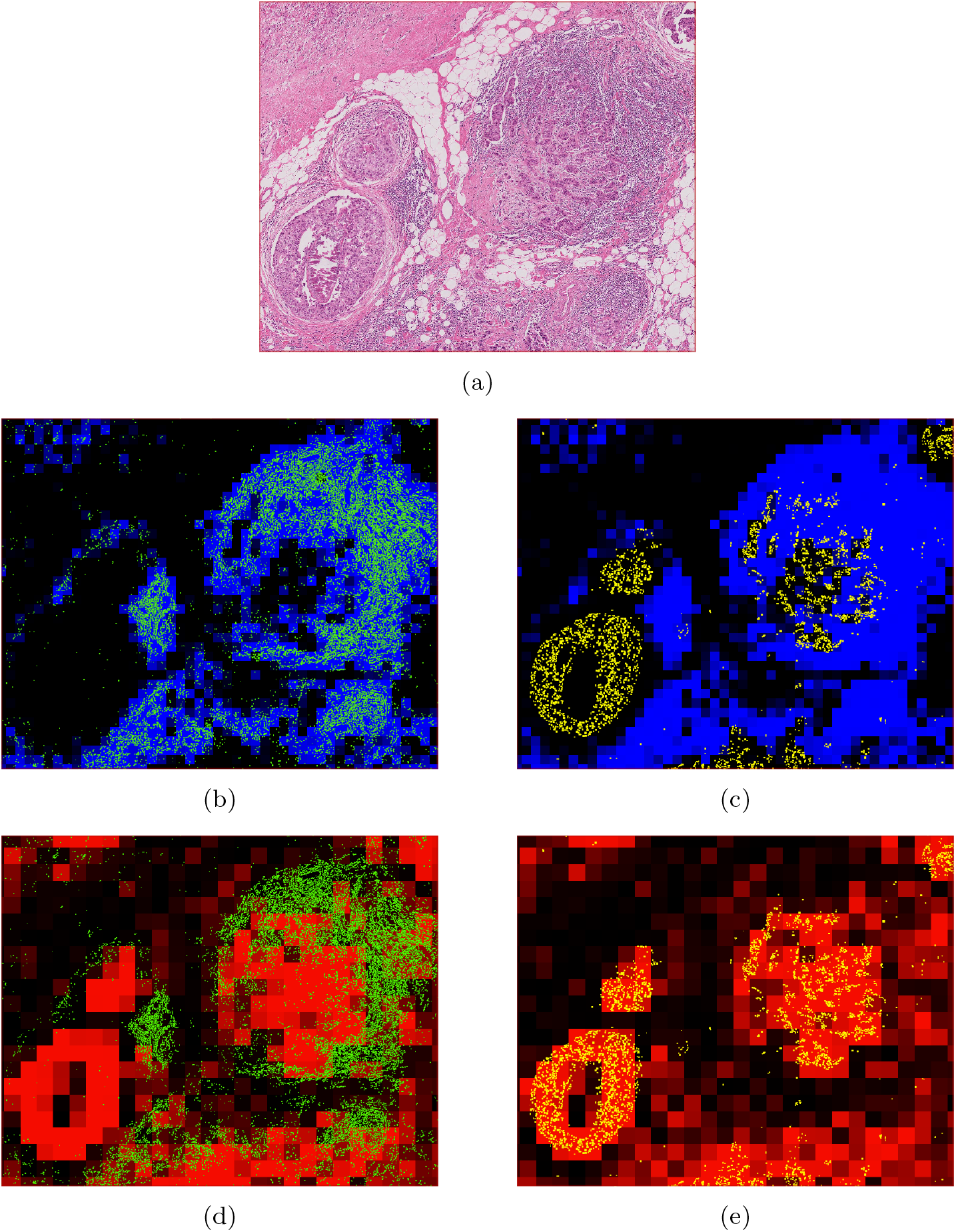
Integrative patch-level and single-cell inference on a breast cancer H&E ROI. **(a)** Representative H&E region of interest. **(b)**–**(e)** The same ROI overlaid with multi-scale predictions from the WSInfer/WSInsight Model Zoo: patch-level heatmaps from `breast-tumor-resnet34.tcga-brca` (tumor probability, red) and `lymphnodes-tiatoolbox-resnet50.patchcamelyon` (lymphocyte aggregation, blue) are co-rendered with cell-level CellViT-SAM-H-x40 detections (inflammatory cells in green; neoplastic epithelial cells in yellow). In particular, panel **(d)** overlays inflammatory-cell detections on the tumor-probability heatmap, so green cells that fall within the red tumor regions correspond to tumor-infiltrating lymphocytes (TILs).

**Fig. 4:**
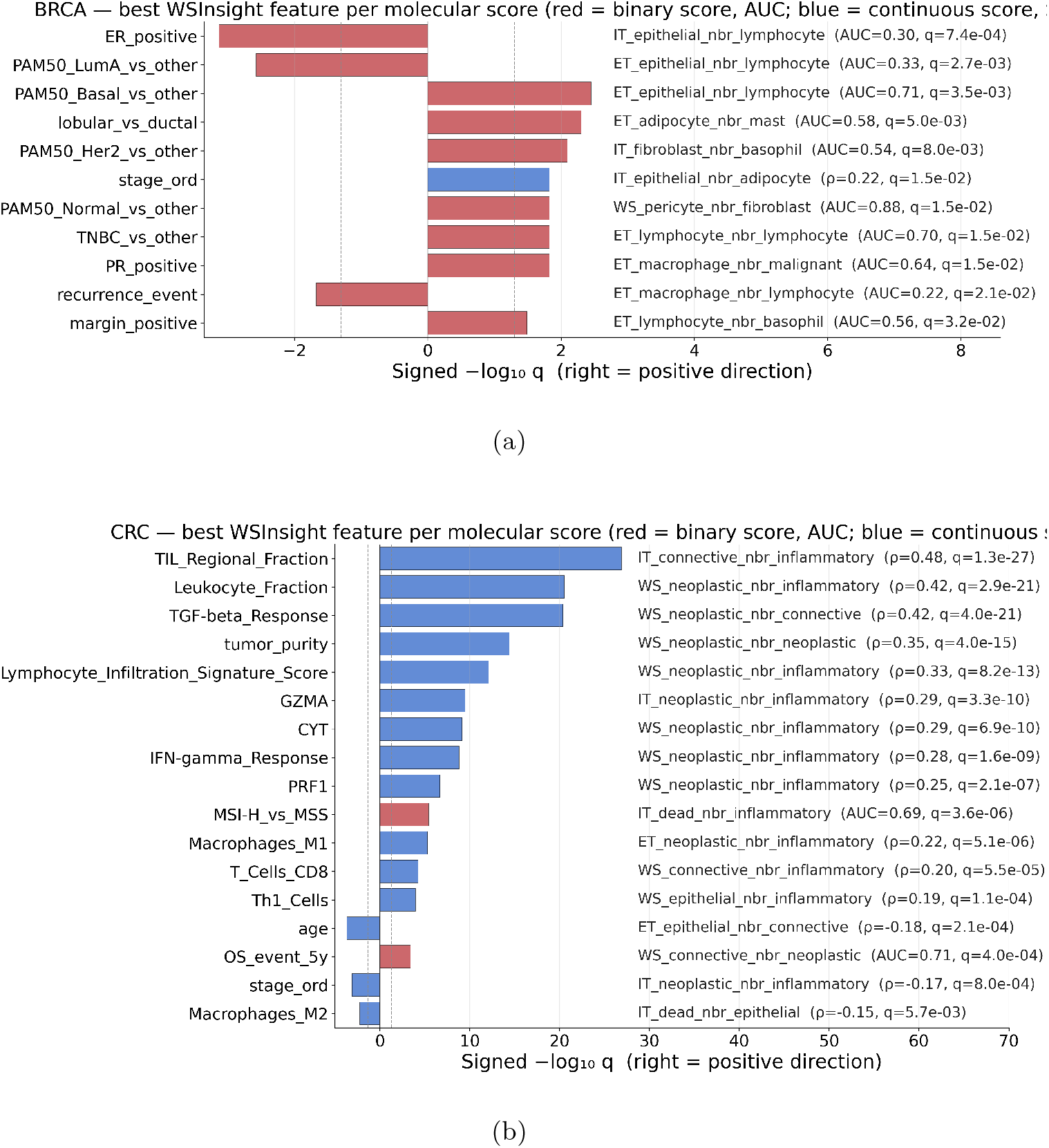
Cross-cohort biomarker landscape produced by a single pipeline invocation. For each molecular, receptor-status, and immune-microenvironment score available in(a) TCGA-BRCA and **(b)** TCGA-CRC, the panel shows the single best WSInsight imaging feature with its detail plot (boxplot+jitter for binary scores with AUC; scatter+linear fit for continuous scores with Spearman *ρ*) and the FDR-adjusted *p*-value. Compartment suffixes use IT (intratumor), ET (extratumor) and WS (whole-slide).

For example, PAM50 Basal slides had a higher TIL ratio than Luminal A slides in TCGA-BRCA (AUC = 0.72 for Basal-vs-other and 0.66 for LumA-vs-other in the cold direction; Fig. 4**(a)**), and MSI-H tumours in TCGA-CRC had higher intratumor TIL density and positive correlations with cytolytic activity, GZMA, and IFN-*γ* response (Fig. 4**(b)**). These are the directions expected from the known Basal/LumA and MSI-H/MSS contrasts. BRCA was processed with the lineage-resolved 10x-Xenium-supervised single-cell head and CRC with the morphology-supervised PanNuke head (see Methods), so cross-cohort comparisons are made at the workflow level rather than at the cell-subtype-feature level.

Indication-specific behaviour was nonetheless preserved: in TCGA-CRC, intratumor TIL density retained an independent protective effect after covariate adjustment, whereas in TCGA-BRCA the same imaging feature did not retain independent prognostic value after adjustment for age, American Joint Committee on Cancer (AJCC) stage, and PAM50 (hazard ratio HR= 0.94, *p* = 0.70), consistent with the known heterogeneity of TIL-based prognosis in hormone receptor-positive disease. Cohort definitions, statistical procedures, and per-endpoint results are reported in Methods (Figs. 6–10). The entire TCGA-scale demonstration was yexecuted as a human-in-the-loop agentic-AI workflow through the WSInsight ClawSight and ClawPyter plugins; the agentic platform, reasoning model, and division of labour between human authors and AI agent are disclosed under *Use of AI tools*.

WSInsight is released as open source under Apache 2.0 (see Code availability). Among the open-source frameworks we compared (Table 1), it is the only one that combines patch- and single-cell H&E inference, direct streaming of whole-slide images from cloud repositories, QuPath and OMERO integration, and an agent-callable interface within one documented workflow. WSInsight addresses a different problem from autonomous discovery systems such as SPARK [15]: rather than generating biological hypotheses, it provides a reproducible WSI analysis runtime that human users, viewer plugins, and AI agents can call to run patch- and single-cell inference, compute neighborhood features, and return standardized outputs to QuPath, OMERO, and downstream statistical workflows. This shared interface is intended to make single-cell whole-slide analysis easier to reproduce across translational and clinical settings.

*Evaluation criteria for Table 1*. A column receives ✓ only if the capability is reachable from the framework’s official entry point without bespoke user glue. **Patch** and **Single-cell** require end-to-end patch-level inference and end-to-end nuclei segmentation with per-cell typing, respectively, on giga-pixel H&E WSIs in a single invocation. **Cloud I/O** requires that input WSIs and outputs can reside on local, S3 or TCGA/GDC storage and are streamed on demand. **QuPath** and **OMERO** require auto-importable per-slide outputs. **Spatial** requires graph-based neighborhood features from the same entry point. **Agent-callable** requires a typed, schema-described tool interface based on a published agent-tool protocol (e.g. MCP).^†^

## Methods

### Model Zoo

WSInsight ships pretrained heads for both patch-level classification and single-cell H&E analysis, and is fully backwards-compatible with the WSInfer Model Zoo so all existing WSInfer patch-level heads run within the WSInsight runtime without modification. The current single-cell collection centres on two complementary families of supervision: (i) morphology-supervised heads (CellViT and HoVer-Net trained on the PanNuke dataset [16]), which provide a tight, biologist-interpretable five-class vocabulary, and (ii) transcriptome-supervised heads (CellViT-SAM-H-x40 fine-tuned with Xenium-derived labels), which expose a richer lineage-resolved vocabulary at the cost of higher per-class noise. A compact architectural comparison is provided in Table 2; the schema and recipe for contributing new heads are documented with the source code (see Code availability).

**Table 2:** Comparison of representative single-cell models [9].

| Method | Architecture and key features | mPQ | bPQ |
| --- | --- | --- | --- |
| **CellViT** [9] | Vision-Transformer encoder with U-Net-style decoder; trained on multi-tissue datasets including PanNuke; supports multi-class nuclear instance segmentation and classification. | 0.4980 | 0.6793 |
| **HoVer-Net** [10] | CNN backbone (typically ResNet50) with dual-branch decoder predicting nuclear masks and horizontal/vertical (HoVer) distance maps for improved instance separation. | 0.4629 | 0.6596 |
| **StarDist-ResNet50** [11, 17] | ResNet50 backbone paired with star-convex polygon representation predicting radial distances along fixed rays for nuclei delineation. | 0.4796 | 0.6692 |

The PanNuke dataset [16] comprises 19,000 H&E image tiles from multiple organs annotated across five morphologic classes (neoplastic, inflammatory, connective, dead, epithelial). PanNuke-trained CellViT and HoVer-Net heads therefore provide a per-class accurate but biologically coarse vocabulary in which every immune cell collapses into a single “inflammatory” bucket. To complement this morphology-only supervision, we additionally assembled a curated collection of 39 public 10x Genomics Xenium datasets spanning 14 tissue sites (Table 3), in which per-cell H&E supervision is derived from paired transcriptomic cluster assignments and therefore preserves immune-lineage detail.

**Table 3:** Public datasets provided by 10x Genomics (under the CC-BY 2.0 license) used for training the WSInsight site-specific single-cell heads.

|  |  |
| --- | --- |
| 1. | Human Bone and Bone Marrow Data with Custom Add-on Panel / Non-diseased Femur Bone |
| 2. | Human Bone and Bone Marrow Data with Custom Add-on Panel / Acute Lymphoid Leukemia Bone Marrow |
| 3. | Human Bone and Bone Marrow Data with Custom Add-on Panel / Non-diseased Bone Marrow |
| 4. | FFPE Human Brain Cancer Data with Human Immuno-Oncology Profiling Panel and Custom Add-on |
| 5. | FFPE Human Breast Cancer with 5K Human Pan Tissue and Pathways Panel plus 100 Custom Genes |
| 6. | FFPE Human Breast using the Entire Sample Area / Replicate 1 |
| 7. | FFPE Human Breast using the Entire Sample Area / Replicate 2 |
| 8. | FFPE Human Breast with Pre-designed Panel / Tissue sample 1 |
| 9. | Xenium FFPE Human Breast with Custom Add-on Panel / Tissue sample 1 (IDC) |
| 10. | FFPE Human Breast with Custom Add-on Panel / Tissue sample 1 |
| 11. | FFPE Human Breast with Pre-designed Panel / Tissue sample 2 |
| 12. | FFPE Human Breast with Custom Add-on Panel / Tissue sample 2 |
| 13. | FFPE Human Cervical Cancer with 5K Human Pan Tissue and Pathways Panel plus 100 Custom Genes |
| 14. | Human Colon Preview Data (Xenium Human Colon Gene Expression Panel) / Cancer / pre-designed + add-on panel |
| 15. | Human Colon Preview Data (Xenium Human Colon Gene Expression Panel) / Non-diseased / pre-designed panel |
| 16. | FFPE Human Colorectal Cancer Data with Human Immuno-Oncology Profiling Panel and Custom Add-on |
| 17. | Human Heart Data with Xenium Human Multi-Tissue and Cancer Panel |
| 18. | Human Kidney Preview Data (Xenium Human Multi-Tissue and Cancer Panel) / Kidney Cancer (PRCC) |
| 19. | Human Kidney Preview Data (Xenium Human Multi-Tissue and Cancer Panel) / Non-diseased Kidney |
| 20. | Xenium In Situ Gene and Protein Expression data for FFPE Human Renal Cell Carcinoma |
| 21. | Human Liver Data with Xenium Human Multi-Tissue and Cancer Panel / Liver cancer |
| 22. | Human Liver Data with Xenium Human Multi-Tissue and Cancer Panel / Non-diseased |
| 23. | Post-Xenium Technical Note: Xenium v1 and Xenium Prime 5K for FFPE Human Lung Cancer / Experiment 2 / Xenium Prime 5K |
| 24. | FFPE Human Lung Cancer Data with Human Immuno-Oncology Profiling Panel and Custom Add-on |
| 25. | Post-Xenium Technical Note: Xenium v1 and Xenium Prime 5K for FFPE Human Lung Cancer / Experiment 1 / Xenium v1 |
| 26. | Preview Data: FFPE Human Lung Cancer with Xenium Multimodal Cell Segmentation |
| 27. | Preview Data: FFPE Human Lymph Node with 5K Pan Tissue and Pathways Panel |
| 28. | FFPE Human Ovarian Cancer with 5K Human Pan Tissue and Pathways Panel plus 100 Custom Genes |
| 29. | FFPE Human Ovarian Cancer Data with Human Immuno-Oncology Profiling Panel and Custom Add-on |
| 30. | Pancreatic Cancer with Xenium Human Multi-Tissue and Cancer Panel |
| 31. | FFPE Human Pancreatic Ductal Adenocarcinoma Data with Human Immuno-Oncology Profiling Panel |
| 32. | FFPE Human Pancreas with Xenium Multimodal Cell Segmentation |
| 33. | Preview Data: FFPE Human Prostate Adenocarcinoma with 5K Human Pan Tissue and Pathways Panel |
| 34. | Preview Data: FFPE Human Skin Primary Dermal Melanoma with 5K Human Pan Tissue and Pathways Panel |
| 35. | Human Skin Preview Data (Xenium Human Skin Gene Expression Panel) |
| 36. | Human Skin Data with Xenium Human Multi-Tissue and Cancer Panel / Sample 1 |
| 37. | Human Skin Data with Xenium Human Multi-Tissue and Cancer Panel / Sample 2 |
| 38. | Human Skin Preview Data (Xenium Human Skin Gene Expression Panel with Custom Add-On) |
| 39. | Human Tonsil Data with Xenium Human Multi-Tissue and Cancer Panel / Follicular lymphoid hyperplasia |
| 40. | Human Tonsil Data with Xenium Human Multi-Tissue and Cancer Panel / Reactive follicular hyperplasia |

Per-cell training labels were derived from paired Xenium transcriptomic cluster assignments registered to H&E coordinates in QuPath and QuST [18], and reviewed for QC by an experienced biologist before being used to fine-tune the breast and colorectal CellViT-SAM-H-x40 heads. Because supervision is transcriptomic rather than morphologic, the resulting heads are biologically richer (typically 11–15 site-specific classes spanning lineage-resolved immune, stromal, and parenchymal phenotypes; Table 4) but per-class noisier than the PanNuke head, and a third party with their own Xenium-paired cohort can produce a site-specific head through the same end-to-end workflow (see Code availability).

**Table 4:** Available single-cell annotations based on 10x Genomics public datasets.

| Tissue site | Labels | Training datasets<br>(see Table 3) |
| --- | --- | --- |
| bone (incl. bone marrow) | adipocyte, chondrocyte, endothelial, fibroblast_like, hematologic_blast, lymphocyte, macrophage_like, malignant_epithelial, mast_cell, neutrophil, osteoblast, osteocyte, pericyte | 1, 2, 3 |
| brain | fibroblast_like, glial, lymphocyte, macrophage_like, neutrophil, pericyte | 4 |
| breast | adipocyte, endothelial, epithelial, fibroblast_like, lymphocyte, macrophage_like, malignant_epithelial, mast_cell, pericyte, plasma_cell | 5, 6, 7, 8, 9, 10, 11, 12 |
| cervix | basaloid_progenitor, endothelial, epithelial, fibroblast_like, lymphocyte, macrophage_like, malignant_epithelial, pericyte, smooth_muscle | 13 |
| colorectal | basaloid_progenitor, basophil, endothelial, eosinophil, epithelial, fibroblast_like, hematologic_blast, lymphocyte, macrophage_like, malignant_epithelial, mast_cell, neuron, neutrophil, pericyte, plasma_cell | 14, 15, 16 |
| heart | basophil, cardiomyocyte, endothelial, fibroblast_like, lymphocyte, macrophage_like | 17 |
| kidney | endothelial, epithelial, fibroblast_like, lymphocyte, macrophage_like, malignant_epithelial, mesangial, neutrophil, pericyte, perivascular, plasma_cell | 18, 19, 20 |
| liver | endothelial, epithelial, fibroblast_like, lymphocyte, macrophage_like, malignant_epithelial, pericyte, plasma_cell | 21, 22 |
| lung | basophil, endothelial, epithelial, fibroblast_like, hematologic_blast, lymphocyte, macrophage_like, malignant_epithelial, mast_cell, pericyte, plasma_cell, smooth_muscle | 23, 24, 25, 26 |
| lymph node | endothelial, fibroblast_like, hematologic_blast, lymphocyte, macrophage_like, mast_cell, pericyte, plasma_cell | 27 |
| pancreas | endothelial, epithelial, fibroblast_like, lymphocyte, macrophage_like, malignant_epithelial, mast_cell, neuroendocrine, neutrophil, pericyte, plasma_cell | 30, 31, 32 |
| prostate | endothelial, epithelial, fibroblast_like, lymphocyte, macrophage_like, malignant_epithelial, neutrophil, pericyte, plasma_cell, smooth_muscle | 33 |
| skin | basophil, endothelial, epithelial, fibroblast_like, lymphocyte, macrophage_like, malignant_epithelial, malignant_melanocytic, mast_cell, melanocyte, neuroendocrine, pericyte, plasma_cell | 34, 35, 36, 37, 38 |
| tonsil | basophil, endothelial, fibroblast_like, hematologic_blast, lymphocyte, macrophage_like, mast_cell, pericyte, plasma_cell | 39, 40 |
| pan-tissue | neoplastic, epithelial, inflammatory, connective, background | all of above |

For the TCGA cohorts in this paper, BRCA was analysed with the lineage-resolved 10x-Xenium-supervised CellViT-SAM-H-x40 head (11 classes) and CRC with the H&E-phenotype-supervised PanNuke head; further site-specific Xenium-supervised heads (Table 4) are added as they are trained. Tiles were exported at 0.25 *µ*m/px. Per-slide Xenium-to-H&E alignment QC is reported in Table 5: across the 8 breast slides, 4,901,986 of 4,926,291 Xenium cells were retained after QuST registration (99.51%). The row-normalised validation confusion matrix for the breast head (Fig. 5**(d)**) is dominated by a strong diagonal; off-diagonal mass is most likely due to H&E morphology ambiguity (e.g. plasma cell →lymphocyte; basophil →lymphocyte), residual class imbalance, and residual transcriptomic-to-morphologic label uncertainty rather than registration noise. Per-class precision, recall and F1 for the breast head are reported with the source code (see Code availability). Lineage-resolved BRCA features are correspondingly framed as exploratory; we do not directly compare them to PanNuke-vocabulary CRC features at the cell-subtype level.

**Table 5:** Per-slide Xenium-to-H&E alignment QC for the 8 training slides used for the breast CellViT-SAM-H-x40 classifier head. Cells were rejected if (i) no cluster assignment in the provided Xenium datasets, (ii) cluster id absent from the per-tissue mapping, or (iii) cell-type string not in the harmonised label map.

| Tissue | Slide stem | Xenium cells | Matched | % |
| --- | --- | --- | --- | --- |
| breast | Xenium_Prime_Breast_Cancer_FFPE | 699,110 | 682,066 | 97.56 |
| breast | Xenium_V1_FFPE_Human_Breast_IDC_Big_1 | 892,966 | 888,925 | 99.55 |
| breast | Xenium_V1_FFPE_Human_Breast_IDC_Big_2 | 885,523 | 882,692 | 99.68 |
| breast | Xenium_V1_FFPE_Human_Breast_IDC_With_Addon<br>(unaligned) | 574,527 | 574,426 | 99.98 |
| breast | Xenium_V1_FFPE_Human_Breast_ILC_With_Addon<br>(unaligned) | 365,604 | 365,592 | 100.00 |
| breast | Xenium_V1_FFPE_Human_Breast_IDC | 574,852 | 574,692 | 99.97 |
| breast | Xenium_V1_FFPE_Human_Breast_ILC | 356,746 | 356,730 | 100.00 |
| breast | Xenium_V1_FFPE_Human_Breast_IDC_With_Addon | 576,963 | 576,863 | 99.98 |
| **Breast subtotal (8 slides)** |  | **4,926,291** | **4,901,986** | **99.51** |

**Fig. 5:**
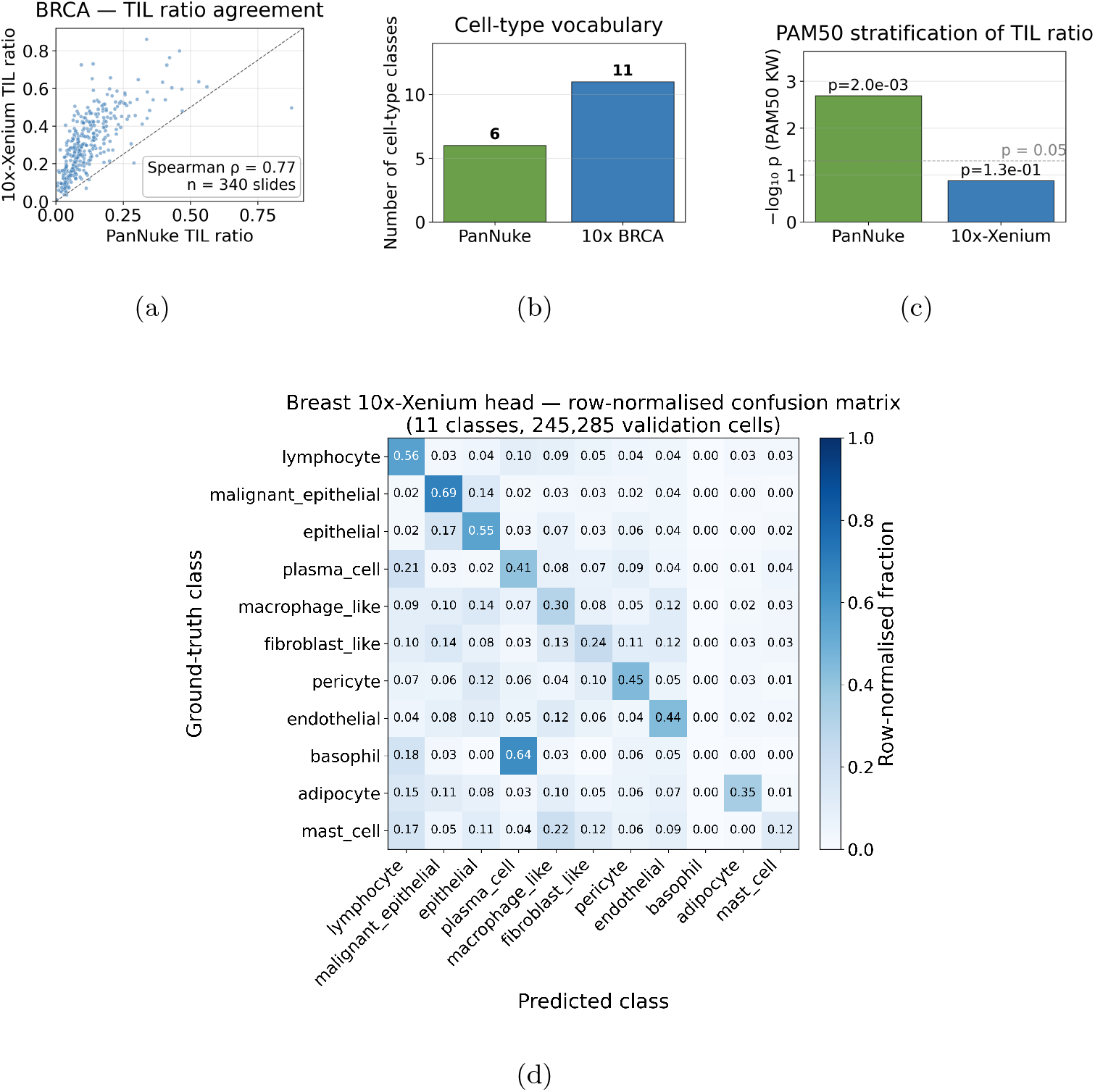
PanNuke vs. 10x-Xenium BRCA head on TCGA-BRCA. PanNuke (H&E phenotype-supervised, 5 classes; the head used for all TCGA-CRC analyses in this paper) vs. the 10x-Xenium-supervised BRCA head (11 classes; the head used for all TCGA-BRCA analyses in this paper). **(a)** Per-slide TIL-ratio agreement (Spearman *ρ* = 0.77, *n* = 340 slides). **(b)** Class vocabulary of each head. **(c)** PAM50 stratification of per-patient TIL ratio (−log_10_ *p, n* = 283): PanNuke *p* = 2.0 ×10^−3^ vs. 10x-Xenium *p* = 1.3× 10^−1^. The higher-vocabulary head’s added value lies in the downstream lineage-specific features it exposes rather than in this single TIL summary. **(d)** Row-normalised validation confusion matrix for the breast 10x-Xenium CellViT-SAM-H-x40 head (11 classes, 8 Xenium slides); rows indicate ground-truth cell type and columns indicate predicted cell type. Per-cell Xenium-to-H&E matched rate after QuST registration was 99.51% (Table 5); the off-diagonal mass likely reflects a combination of H&E morphology ambiguity, class imbalance, and residual transcriptomic-to-morphologic label uncertainty rather than registration label noise alone.

**Fig. 6:**
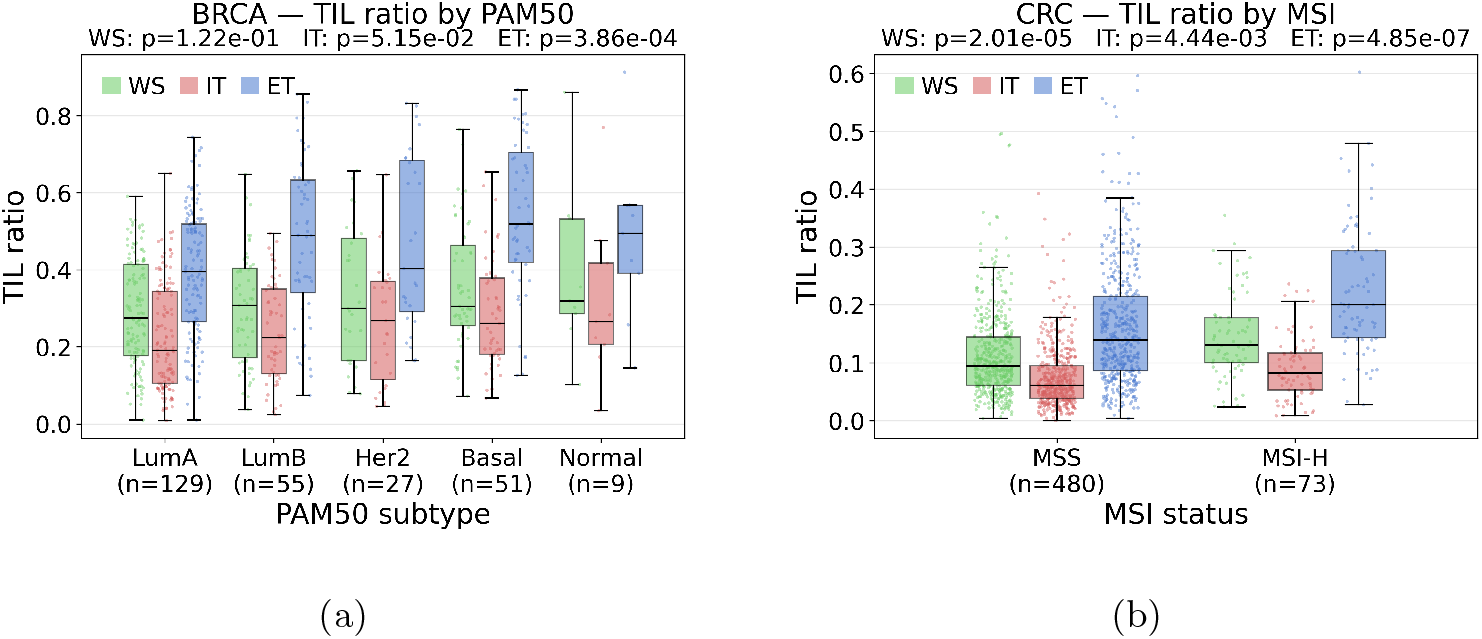
TIL ratio (lymphocyte fraction) across molecular subtype. **(a)** TCGA-BRCA stratified by PAM50 (*n* = 301); per-stratum significance reported by Kruskal–Wallis.(b) TCGA-CRC stratified by MSI status (*n* = 310); per-stratum significance reported by Mann–Whitney *U*. In both panels the three strata per subtype—IT (intratumor, red), ET (extratumor, blue), WS (whole-slide, green)—are shown as adjacent boxplots within each group, and jittered points are individual patients.

**Fig. 7:**
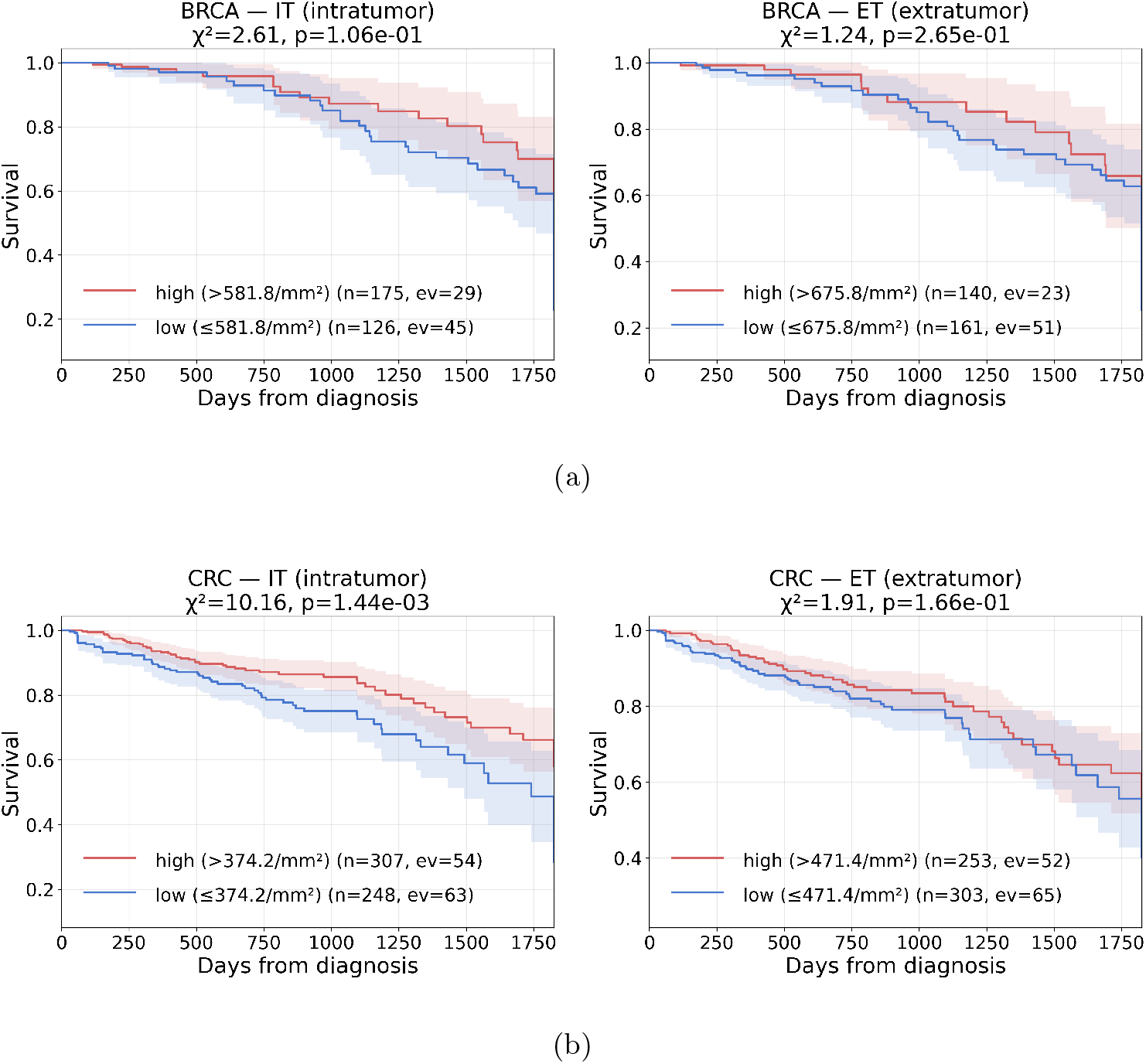
Exploratory survival panels for intratumor (IT) and extratumor (ET) TIL density at 5-y OS, with optimal log-rank cutoff (10–90th percentile, ≥10 events per arm) and Greenwood 95% ribbons; intended as platform-output sanity checks rather than validated prognostic claims. The whole-slide (WS) stratum is omitted because it is the union of IT and ET and therefore carries no information beyond the two compartments. **(a)** TCGA-BRCA: trends toward better survival in the high-TIL arm are observed in both compartments but neither reaches nominal significance in this exploratory screen (IT *p* = 1.1 ×10^−1^; ET *p* = 2.6 ×10^−1^). **(b)** TCGA-CRC: both compartments separate at nominal significance, with IT slightly stronger than ET (IT *p* = 2.2× 10^−2^; ET *p* = 2.9× 10^−2^). Post-resection systemic therapy is heterogeneous and incompletely recorded in TCGA [19] and is therefore not adjusted for in these curves.

**Fig. 8:**
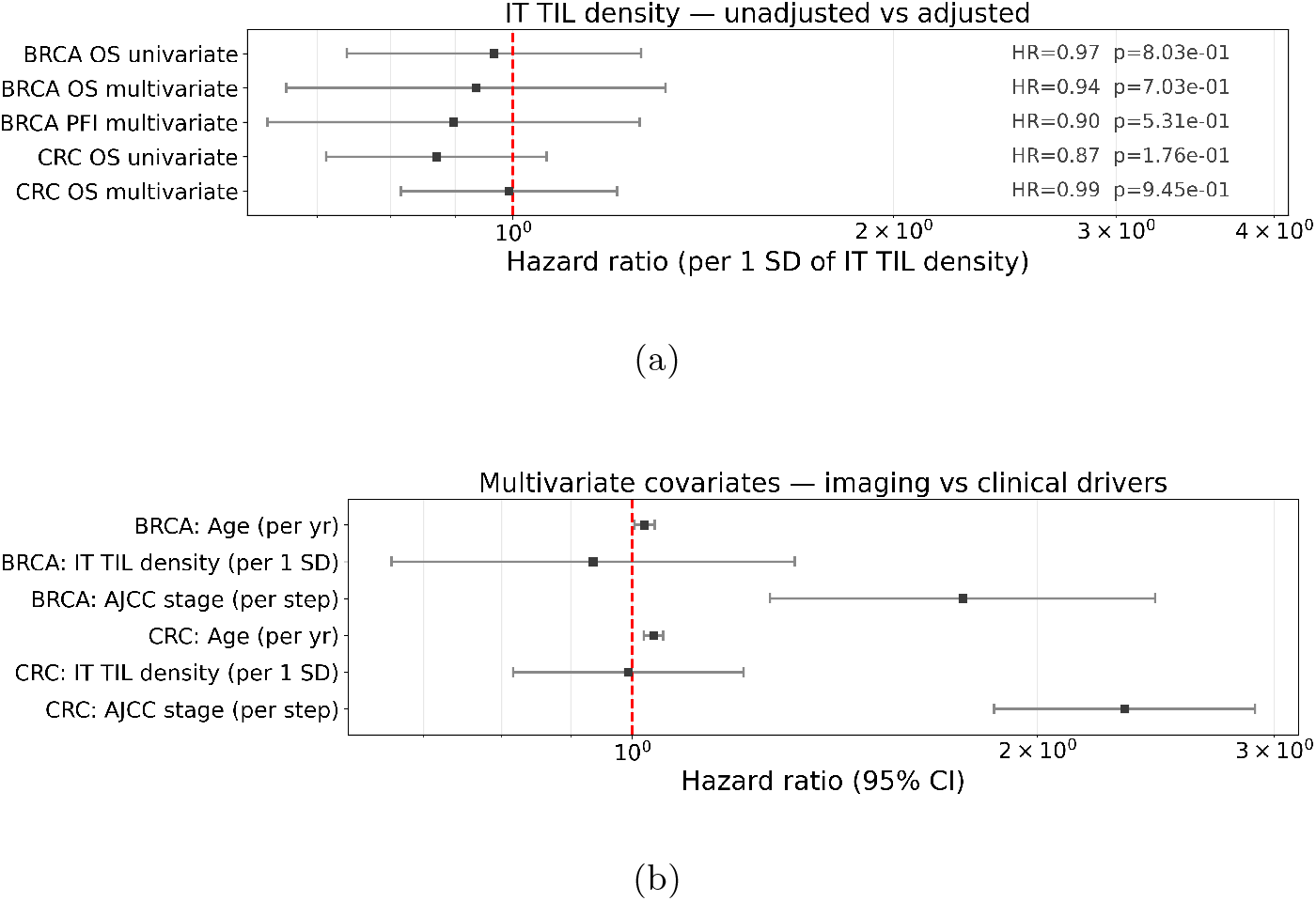
Confounder-adjusted Cox analysis of intratumor TIL density (HR per 1 SD on the continuous, *z*-scored feature). **(a)** Imaging-feature HR across five models— unadjusted vs. adjusted on age + AJCC stage + PAM50 (BRCA OS, BRCA PFI) or + MSI (CRC OS). **(b)** Imaging-vs.-clinical comparison in the multivariate models, showing that AJCC stage carries the dominant prognostic weight in CRC. Confidence intervals are 95%.

**Fig. 9:**
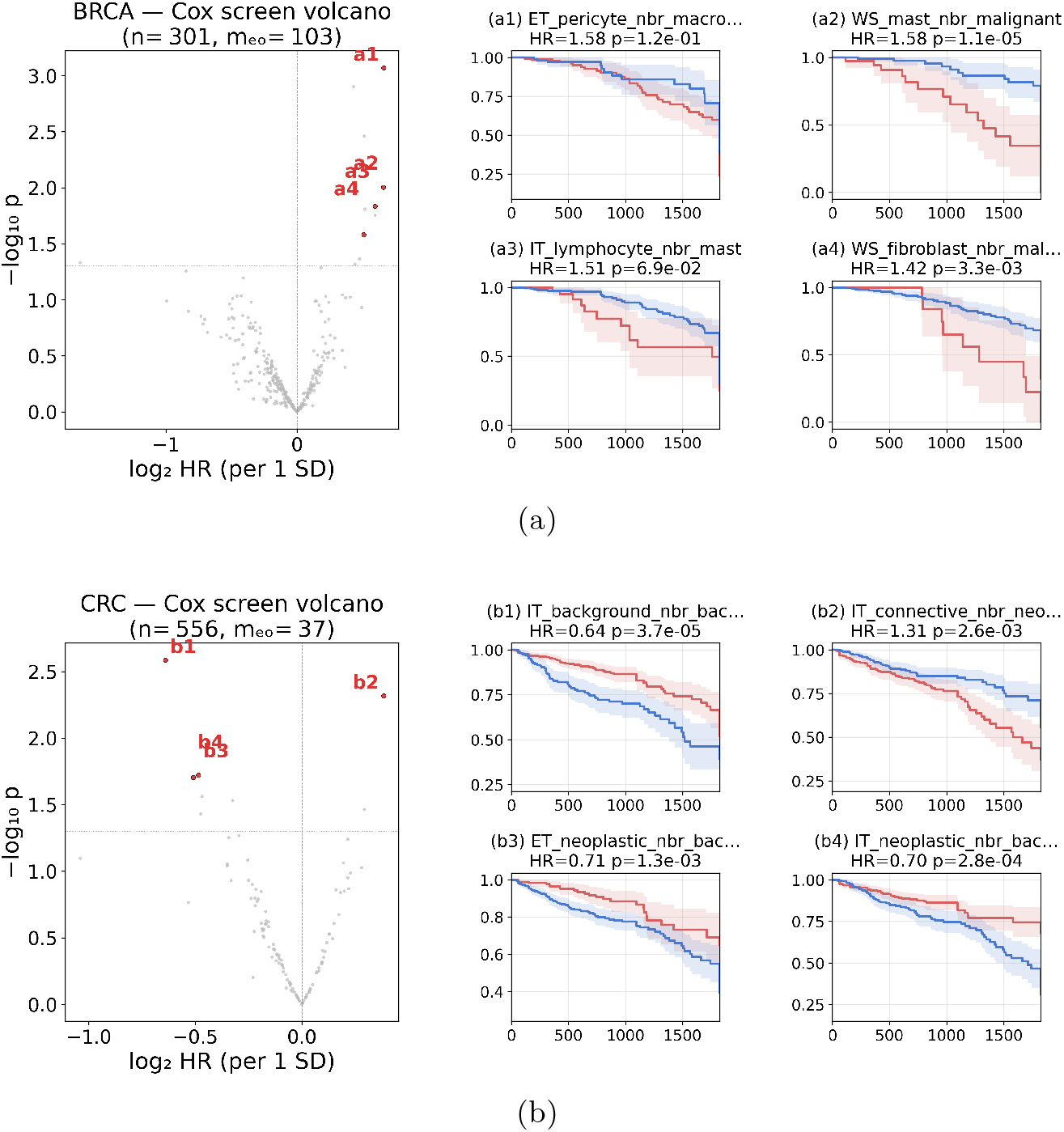
Whole-panel Cox screen (volcano plots). **(a)** BRCA volcano plot of log_2_ HR per 1 SD against −log_10_ *p* for every WSInsight feature; the four lowest-Cox-*p* features that admit a valid log-rank dichotomization (10–90th percentile, ≥10 events per arm, ≥30 patients with non-missing values) are coloured and named so they can be matched to the KM panels at the optimal cutoff shown to the right. **(b)** CRC equivalent. In both panels the horizontal dashed line marks nominal *p* = 0.05 and the vertical dashed line marks log_2_ HR = 0 (no effect). The corresponding top-10 Cox-ranked features per cohort, with feature-cluster-corrected BH-FDR *q*-values, are shown in Fig. 10.

**Fig. 10:**
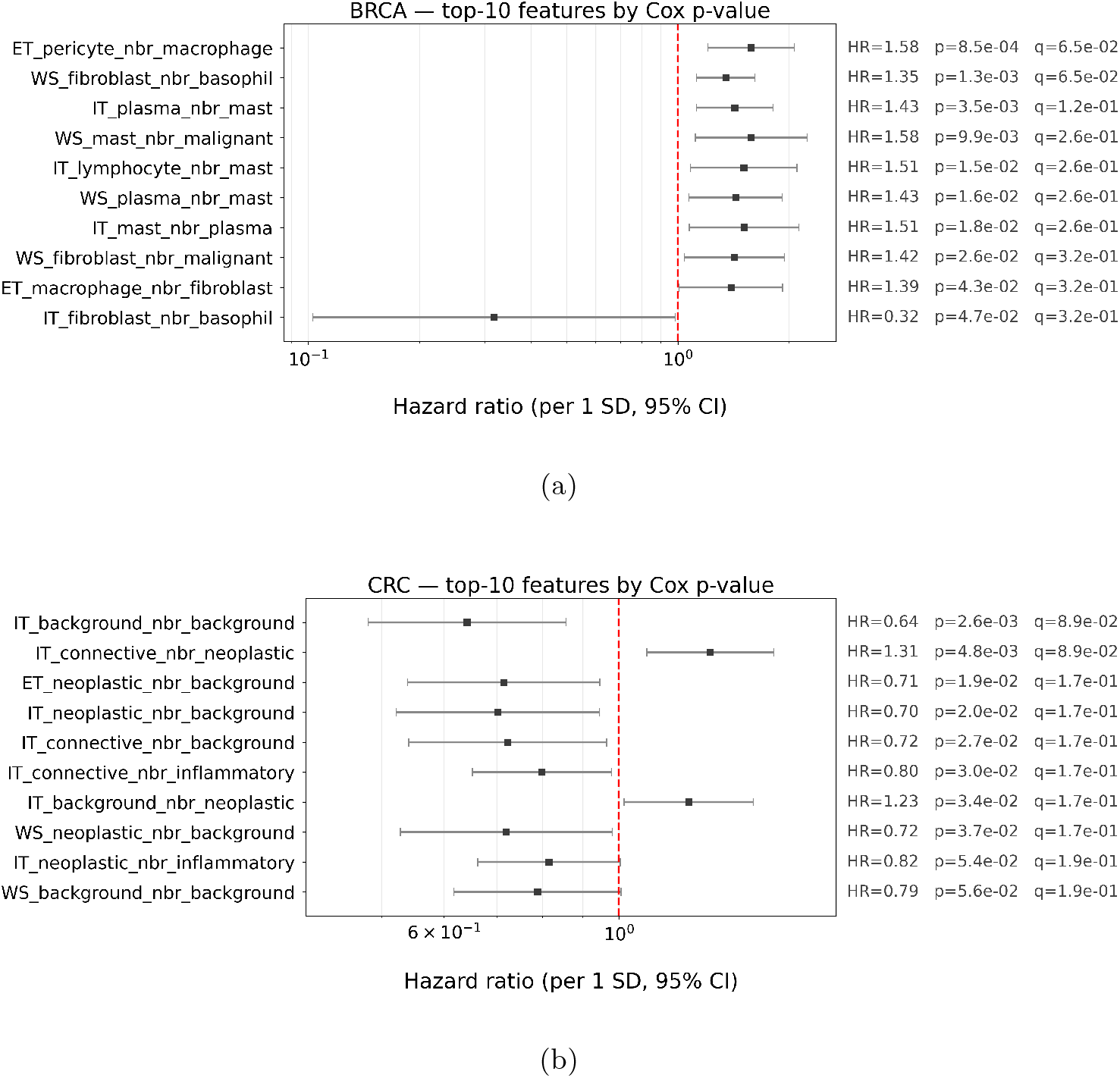
Top-10 WSInsight features ranked by Cox *p* from the whole-panel screen of Fig. 9, displayed as forest plots. Each row shows the per-1 SD hazard ratio (point estimate) with its 95% confidence interval as whiskers; the *x*-axis is hazard ratio on a log_10_ scale and the red dashed vertical line marks HR = 1 (no effect). Per-feature HR, raw Cox *p*, and feature-cluster-corrected BH-FDR *q*-values are annotated to the right of each row; the BH denominators are the effective numbers of independent feature clusters at Spearman *ρ* ≥ 0.7 (*m*_eff_ = 103 BRCA, 40 CRC). (a) TCGA-BRCA. (b) TCGA-CRC.

### Comparison of PanNuke and 10x-Xenium heads

Among the public 10x Genomics Xenium releases, breast is the largest single-tissue cohort (eight slides), making it the natural setting in which to compare a transcriptome-supervised head against the H&E phenotype-supervised PanNuke head. We therefore ran both heads on TCGA-BRCA (*n* = 340 slides; *n* = 283 patients with PAM50 calls) and compared two readouts: the per-slide TIL ratio and its per-patient PAM50 stratification.

On the per-slide TIL ratio, the two heads agree at Spearman *ρ* = 0.77 (*n* = 340; Fig. 5**(a)**), while their class vocabularies differ by ∼2× (6 vs. 11 classes; Fig. 5**(b)**). On the coarse PAM50 stratification of TIL ratio, PanNuke yields Kruskal–Wallis *p* = 2.0×10^−3^ and the 10x-Xenium head yields *p* = 1.3×10^−1^ (*n* = 283; Fig. 5**(c)**).

### Inference runtime

We benchmarked the single-cell inference runtime on a server equipped with eight NVIDIA H100 graphics processing units (GPUs), using the breast Xenium-supervised CellViT-SAM-H-x40 head. The TCGA-BRCA cohort was sharded into eight parallel subsets, with one worker pinned to each GPU, processing 340 WSIs in total (∼41– 43 WSIs per worker). The full inference job finished in approximately 58 h 25 min of wall-clock time, corresponding to an aggregate throughput of ∼5.8 WSIs per hour, or approximately 82 min per WSI when normalized to a single-GPU system.

### Spatial analysis

WSInsight provides two graph-based spatial-analysis modules that operate on the per-cell detections produced by the inference runtime to characterize the tumour microenvironment: (i) object-to-region spatial registration, which links single-cell detections to patch-level predictions, and (ii) neighborhood-composition profiling (ncomp), which summarises the local cellular context of every cell.

#### Object-to-region spatial registration

Each cell’s centroid is intersected with patch-level region predictions, and region-level attributes (e.g. tumour probability, tissue class) are propagated to the matching cell record.

#### Neighborhood composition ncomp)

Given *N* cells with centroids 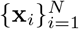 and cell-type assignments *c*_*i*_ ∈ {1, …, *C*}, the module proceeds in three steps. First, a Delaunay triangulation is computed over all cell centroids; edges longer than a configurable Euclidean distance threshold *d*_max_ (default 25 *µ*m, approximately one cell diameter at 0.25 *µ*m/px in standard FFPE H&E and consistent with Delaunay-based cellular-neighborhood methods in spatial biology) are pruned so that connectivity reflects physical contact or near-contact. The resulting graph is encoded as a sparse symmetric adjacency matrix **A** ∈ {0, 1}^*N* ×*N*^ with **A**_*ij*_ = 1 iff cells *i* and *j* are connected and ∥**x**_*i*_ − **x**_*j*_∥ ≤ *d*_max_. Second, the *K*-hop neighborhood of every cell is obtained from the augmented adjacency **Ã** = **A** + **I** via sparse matrix exponentiation, **M** = **Ã** ^*K*^, where **M**_*ij*_ > 0 indicates that cell *j* is reachable from cell *i* within *K* hops; the default *K*=2 balances locality and statistical stability. Binarising **M** yields the *K*-hop neighbour set 𝒩 (*i*) = { *j* : **M**_*ij*_ > 0, *j* ≠*i* }. Third, within each 𝒩 (*i*) the module tallies both the count and the proportion of every annotated cell type, *n*_*i,c*_ =∑ _*j*∈ 𝒩 (*i*)_ **1**[*c*_*j*_ = *c*] and *p*_*i,c*_ = *n*_*i,c*_/| 𝒩 (*i*)|, producing a compact 2*C*-dimensional descriptor (per-class counts concatenated with per-class proportions) of each cell’s spatial context.

#### Propagation of classifier error into ncomp features

Because *c*_*i*_ is the head’s predicted label, classifier confusion (Fig. 5**(d)**) propagates linearly into the per-cell composition tallies 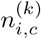. Users of the lineage-resolved BRCA features should therefore interpret `ncomp_ *_ lymphocyte` bins as a lymphocyte-plus-plasma readout rather than a pure lymphocyte readout, and `ncomp_*_fibroblast_like` bins as a stromal-plus-perivascular readout. Bins keyed on PanNuke classes are unaffected because the PanNuke head’s recall is high and its vocabulary is morphology-tight.

### Use case: end-to-end analysis on TCGA cohorts

Both cohorts were restricted to pretreatment, primary-tumour FFPE diagnostic (“DX”) H&E slides retrieved from the NCI GDC; frozen-section slides and post-treatment material were excluded. After this filter, the cohorts comprised *n* = 340 slides (TCGA-BRCA) and *n* = 310 patients (TCGA-CRC). For each TCGA-BRCA and TCGA-CRC slide we extracted neighborhood-composition features summarizing the mean 2-hop cell-type composition around every center cell type, plus per-stratum TIL ratio and TIL density, computed in three strata—intratumor (IT), extratumor (ET), and whole-slide (WS)—using the Delaunay graph with edges truncated at 25 *µ*m. Throughout, the TIL ratio of a slide-stratum is the fraction of detected cells classified as lymphocytes within that stratum (dimensionless, in [0, 1]), and the TIL density is the count of detected lymphocytes within the stratum divided by the QC-defined annotated tissue area of that stratum (in cells / mm^2^); both readouts are platform use-case summaries rather than clinical biomarkers. Slide-level features were averaged to the patient level and screened univariately against every available clinical or molecular score per cohort with Benjamini–Hochberg FDR correction. The per-stratum (IT / ET / WS) TIL-ratio distributions across molecular subtype are shown in Fig. 6, with PAM50 stratification in BRCA assessed by Kruskal–Wallis and MSI stratification in CRC by Mann–Whitney *U*. For survival readouts, TIL density was used as the imaging summary and patients were dichotomized at the optimal log-rank cutoff (10–90th percentile, ≥10 events per arm) for 5-year censored OS (Fig. 7). Confounder-adjusted Cox models were fit on the continuous, *z*-scored intratumor TIL density (HR per 1 standard deviation, SD), adjusting for age, AJCC stage, and molecular subtype (PAM50 in BRCA, MSI in CRC); a BRCA progression-free interval (PFI) sensitivity model and a subtype-stratified Cox at the optimal cutoff are reported alongside (Fig. 8). The whole-panel univariate Cox screen (Fig. 9) was run across all 369 BRCA and 114 CRC WSInsight features (continuous, HR per 1 SD); the effective number of independent tests was estimated by hierarchical clustering of features at Spearman *ρ* ≥0.7 and the cluster count was used as the BH-FDR denominator (*m*_eff_ = 103 BRCA, 40 CRC).

### Interoperability with QuPath, OMERO, and agentic AI

#### QuPath

A bundled QuPath [13] extension lets pathologists run WSInsight inference on the active project without leaving the viewer; per-slide GeoJSON detections and OME-CSV cell tables are imported back into the QuPath project for immediate review.

### OMERO

A companion OMERO module registers the WSInsight OME-CSV cell tables as ROIs on the corresponding image in OMERO [14], with per-object measurements queryable from downstream OMERO workflows.

#### AI agent interface

WSInsight exposes its command-line interface to AI agents through a standards-conformant Model Context Protocol (MCP) server, so an agent and a human user invoke the same commands with the same arguments. The MCP server is reachable from contemporary agentic platforms, including Claude Code [20], OpenClaw [21], and Hermes [22], with companion plugins for OpenClaw and Hermes released alongside WSInsight (see Code availability). Downstream statistical and visualization workflows in this study were authored as Jupyter notebooks and remain the primary working environment for human analysts; to give agentic platforms an equivalent point of access to that environment, we additionally provide ClawPyter, a lightweight bridge that lets agents read, write, and execute notebook cells in the same way a human user would (see Code availability). Because whole-slide inference can take hours on a GPU, long-running steps are issued as asynchronous jobs that an agent can monitor and that respect the same cancellation behaviour as the interactive workflow. A complete agent-driven reproduction of the TCGA-CRC analysis is documented with the source code (see Code availability).

## Ethics statement

This study used only publicly available, de-identified human data from TCGA via the NCI GDC [12], and publicly available 10x Genomics Xenium reference datasets distributed under the Creative Commons Attribution 2.0 license. Informed consent and ethical approval for the original TCGA tissue samples were obtained by the contributing tissue source sites under their respective institutional review board protocols. No new patient data, tissue, or images were generated for this work.

## Data availability

All data used in this study are derived from publicly available resources; no new patient data were generated. The specific resources are summarized in Table 6.

**Table 6:** Public data resources used in this study.

| Resource | Source / URL |
| --- | --- |
| TCGA whole-slide images (BRCA, CRC) | NCI GDC portal [12]<br>https://portal.gdc.cancer.gov |
| TCGA clinical annotations | PanCancer Clinical Data Resource [19] |
| TCGA MSI status calls | Published MSI status table [23] |
| Immune-microenvironment scores | Pan-cancer immune landscape [24]; cytolytic activity per [25] |
| 10x Genomics Xenium reference datasets | 10x Genomics public datasets portal (specific datasets enumerated in Table 3)<br>https://www.10xgenomics.com/datasets |
| Pretrained model weights, WSInfer | Hugging Face Hub<br>https://huggingface.co/kaczmarj |
| Pretrained model weights, WSInsight | Hugging Face Hub<br>https://huggingface.co/huangch |

## Code availability

All WSInsight components are open source under the Apache 2.0 license. The repositories below (Table 7) are organized by their role in the platform. The core repository exposes a self-describing entry point so that the pipeline can be discovered and invoked programmatically by AI agents or scripted workflows; the TCGA-CRC analysis in this paper is reproducible end-to-end through two CLI calls (or the equivalent MCP invocations) and a worked example is documented in the repository.

**Table 7:** WSInsight open-source components and where to obtain them.

| Component | Description / URL |
| --- | --- |
| Core pipeline | Inference runtime, container images, Model Zoo configurations<br>https://github.com/huangch/wsinsight |
| QuPath extension | Pathologist-facing viewer integration<br>https://github.com/huangch/qupath-extension-wsinsight |
| OMERO module | Cell-table registration as ROIs in OMERO<br>https://github.com/huangch/omero-plugin-wsinsight |
| Agent plugins (OpenClaw, Hermes) | MCP-conformant agent surface<br>https://github.com/huangch/clawsight |
| Agent-JupyterLab bridge (ClawPyter) | Notebook read/write/execute bridge for the reproduction notebooks<br>https://github.com/huangch/clawpyter |
| Model-development workflow | Training pipelines, dataset curation, recipes for contributing new heads<br>https://github.com/huangch/wsinsight-model-development |
| Pre-built Docker image | https://hub.docker.com/r/huangchtw/wsinsight |

## Use of AI tools

The TCGA-scale demonstration reported in this paper was executed as a human-in-the-loop agentic-AI workflow. The authors directed AI agents [21, 22], through two WSInsight companion plugins: ClawSight, which exposes the WSInsight CLI as a Model Context Protocol tool surface; and ClawPyter, which lets the agent read, write, and execute JupyterLab notebook cells (see Code availability). AI agents assisted with GDC manifest selection, execution of the WSInsight inference and neighborhood-composition pipeline, and generation of the reproduction notebook used to produce the figures and statistics presented here. The human authors specified the scientific design, reviewed the generated code and outputs, selected and interpreted the analyses, and take full responsibility for all conclusions. AI tools were not listed as authors, in accordance with current journal policy. The cohort-scale results presented here are intended as a platform demonstration of the agent-callable workflow, and should not be interpreted as independent scientific discoveries or clinical biomarker claims; all reported associations are exploratory and require prospective validation outside the scope of this work.

## Acknowledgements

We thank the WSInfer authors and the digital pathology and computational biology communities for open-source resources, and acknowledge TCGA and 10x Genomics for the public datasets that enabled this study. No external funding was received; this work was supported by Pfizer Inc.

## Author contributions

C.H.H. conceived and led the WSInsight design and implementation. O.E.A. contributed model validation and dataset curation. D.F. oversaw oncology use cases and biological interpretation. All authors wrote and approved the manuscript.

